# Extended *in vitro* inactivation of SARS-CoV-2 by titanium dioxide surface coating

**DOI:** 10.1101/2020.12.08.415018

**Authors:** P Mlcochova, A Chadha, T Hesselhoj, F Fraternali, JJ Ramsden, RK Gupta

## Abstract

SARS-CoV-2 transmission occurs via airborne droplets and surface contamination. We show tiles coated with TiO_2_ 120 days previously can inactivate SARS-CoV-2 under ambient indoor lighting with 87% reduction in titres at 1h and complete loss by 5h exposure. TiO_2_ coatings could be an important tool in containing SARS-CoV-2.

## Background

Within 10 months, severe acute respiratory syndrome coronavirus 2 (SARS-CoV-2) has infected 62 million people and has killed over 1.5 million worldwide. Respiratory droplets are believed to be the major vehicle of SARS-COV-2 transmission. Droplets or other body fluids from infected individuals can contaminate surfaces and viable virus has been detected on such surfaces, including surgical masks, for hours, even days depending on different factors including humidity, temperature and type of surface.^1,2,3^ One therefore infers that any external contamination of PPE may last hours or even days.

Recently, there has been a further increase in SARS-CoV-2 cases across Europe and the UK, despite severe mitigation measures following the first wave in the first half of 2020. More worryingly, nosocomial transmission within hospitals is being observed despite universal adoption of wearing face masks, regular testing of staff and patients, and social distancing measures. It may well be that contamination of surfaces is now disproportionately contributing to transmission.^4,5^

Traditional forms of decontamination (such as alcohol-based sprays, quaternary ammonium compounds, and sodium hypochlorite and other chlorine-based compounds) require repeated applications. Photocatalytic surfaces, on the other hand, permanently oxidize, inactivate and destroy microorganisms under normal ambient lighting conditions.^6^ A recent hospital study of titanium dioxide-coated surfaces demonstrated progressive lowering of the bacterial bioburden.^7^ Moreover, the radicals are not considered to induce antimicrobial resistance.^8^ TiO_2_ is especially attractive because it is considered nontoxic to humans: titanium, coated with its oxide, is the most widely used material for implants.^9^ TiO_2_ is also exceedingly stable, unlike other photocatalysts such as zinc oxide and tungsten trioxide. Illumination of TiO_2_ generates highly oxidizing free radicals that are known to have bactericidal and antiviral action against influenza and rotavirus.^10,11,12^ SARS-CoV-2 has not hitherto been investigated.

## Methods and materials

### Cell lines

293T and Vero E6 cells were cultured in DMEM complete (DMEM supplemented with 100 U/ml penicillin, 0.1 mg/ml streptomycin, and 10% fetal calf serum). Vero E6 were a gift from Prof. Ian Goodfellow. 293T cells were obtained from ATCC. ACE-2/TMPRSS2-expressing 293T cells were generated as described previously.^13^

### Pseudotyped virus

SARS-CoV-2 Spike pseudotyped HIV-1 luciferase particles were produced by transfection of 293T cells with pCAGGS-SARS-CoV-2 spike, p8.91HIV-1 gag-pol expression and pCSFLW (expressing the firefly luciferase reporter gene with the HIV-1 packaging signal) as previously described.^15^ Viral supernatant was collected at 48 and 72 h after transfection, filtered through a 0.45 μm filter and stored at −80 °C. The 50% tissue culture infectious dose (TCID_50_) of SARS-CoV-2 pseudovirus was determined using the Steady-Glo luciferase assay system (Promega).

### Viral isolate

Live SARS-CoV-2 (SARS-CoV-2/human/Liverpool/REMRQ0001/2020) used in this study was isolated by Lance Turtle (University of Liverpool), David Matthews and Andrew Davidson (University of Bristol). A SARS-CoV-2 virus stock was produced by infecting Vero E6 cells at MOI 0.01. Culture supernatant was collected 48 h post-infection. The titre of the stock was determined by adding tenfold serial dilutions of virus onto Vero E6 cells. 24 h post-infection cells were fixed, stained for nucleocapsid protein (anti-Nucleocapsid antibody, MA5-36086, ThermoFisher) and %infection determined by flow cytometry. SARS-CoV-2 virus titres were determined as infectious units per ml (IU/ml) as follows: (% infected cells) × (total number of cells) × (dilution factor) / volume of inoculum added to cells.

### Surfaces and illumination

Virus stability was evaluated on the following surfaces: sterile untreated Sterilin standard Petri dish; TiO_2_- and TiO_2_–Ag (Ti:Ag atomic ratio 1:0.04)-coated 45 × 45 mm ceramic tiles (Invisi Smart Technologies UK Ltd). The coatings are transparent and colourless and therefore invisible to the human eye. After coating the tiles were stored for 2–4 months before use. Surfaces were exposed (610 lx, ambient laboratory light) for 1 h before the start of each experiment to ensure a steady state of radical generation. The same light was used during virus exposure, during which relative humidity was approximately 65% and temperature 21 °C.

### Surface inoculation and sampling

#### SARS-CoV-2 spike pseudotyped virus inactivation

Tile surfaces were inoculated with 10^5^ RLU of SARS-CoV-2 spike pseudotyped HIV-1 luciferase virus at time *t* = 0 and illuminated for up to 6 h. At intervals virus was recovered from surfaces with DMEM complete followed by infection of ACE-2/TMPRSS2-expressing 293T cells. Luminescence was measured using Steady-Glo Luciferase assay system (Promega) 48 h post-infection.

#### SARS-CoV-2 live virus inactivation

6×10^6^ IU/ml of SARS-CoV-2 virus was added onto the surface of the tiles at a dosage of 2 μl over 5 × 5 mm. After illuminating for 0–300 min virus was recovered from surfaces with DMEM complete followed by infection of Vero E6 cells. % of infected cells was determined by flow cytometry detecting SARS-CoV-2 nucleocapsid protein 24 h post-infection.

#### Kinetic analysis of inactivation

The main challenge is that laboratory inactivation experiments are necessarily carried out with large numbers of viruses, with which the inactivating material is brought into contact at the beginning of the experiment, and the decay of the entire virus population is measured.^14^ What is of practical interest in the scenario of a coating designed to keep surfaces (e.g., in a hospital) free of viral (and bacterial) bioburden is how quickly an individual virus is inactivated. According to analysis of previously reported results for influenza virus inactivation,^11^ the kinetics fit a convective diffusion transport model even in the absence of mechanical agitation, most likely due to almost inevitable thermal gradients.^14^ The concentration of survivors is thereby predicted to follow a so-called exponential decrease, and plotting the logarithm of the number of survivors v. time should give a straight line, the slope of which is –*k*, the inactivation rate coefficient. The value of *k* can then be compared with the transport-limited fastest possible rate calculated from the size of the virus.^14^

## Results

After 1 h illumination the pseudotyped viral titre was decreased by four orders of magnitude (Figure 1A). There was no significant difference between the TiO_2_ and TiO_2_–Ag coatings. Light alone had no significant effect on viral viability.

**Figure:**
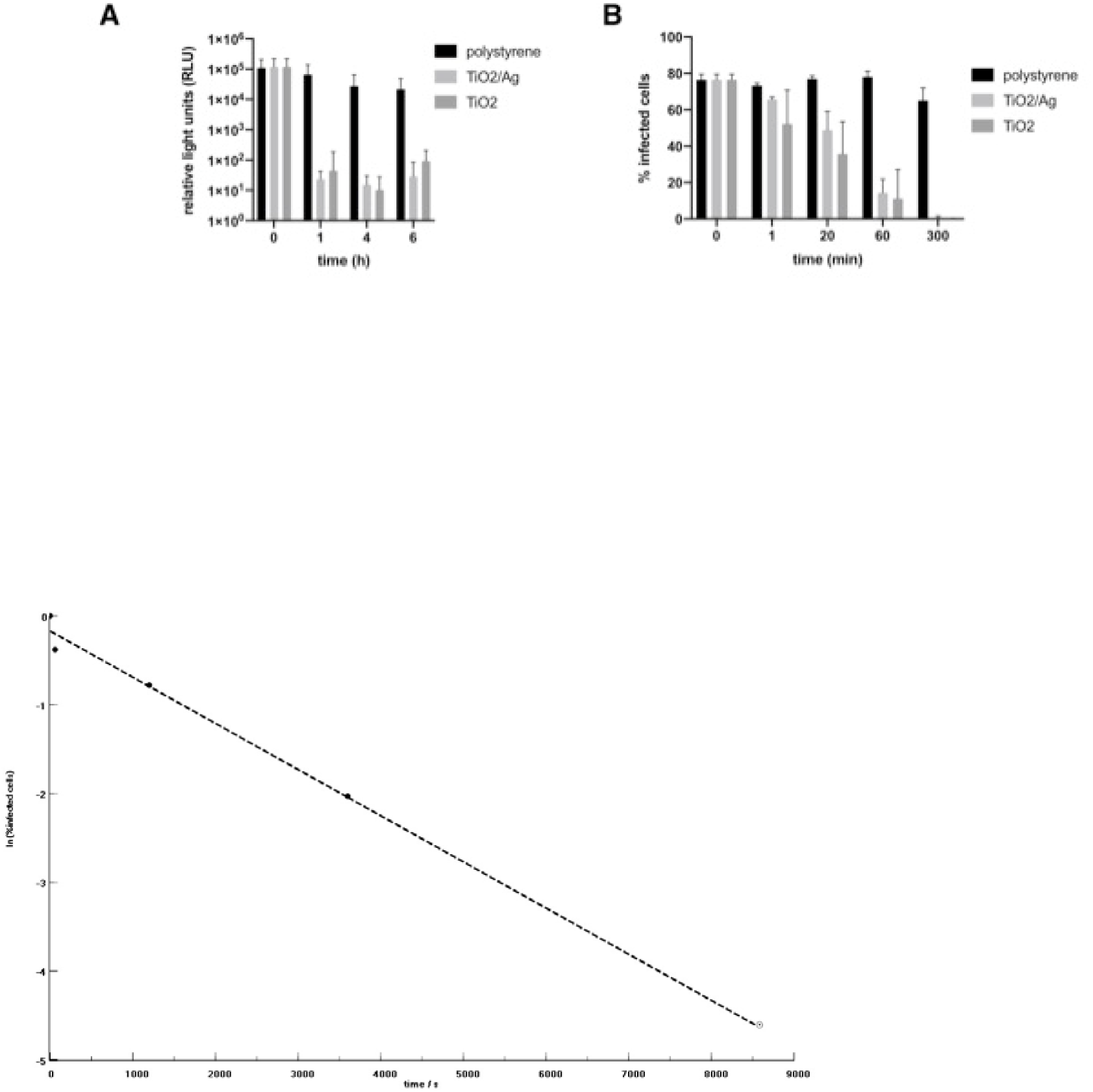
Effect of lnvisiSmart TiO2/Ag and TiO2 tiles on **(A)** SARS-CoV-2 Spike pseudotype viral infection and **(B)** SARS-CoV-2 SARS-CoV-2 isolate REMRQ0001/Human/2020/Liverpool. (C) Plot of the data from (B) as ln(%infected cells) v. time. Dashed line, linear regression using the first 4 data points, the slope of which is *k*. The 5 h titre **(zero)** cannot be plotted as its logarithm but extrapolating the regression line to the estimated virus detection limit (1%) suggests that complete inactivation was effectively reached at 2 h 20 min (open circle).

Next, we tested the ability of the coated tiles to inhibit fully infectious live virus. Coated and uncoated surfaces were exposed to SARS-CoV-2. Virus was harvested at the times indicated and used to infect Vero E6 target cells. SARS-CoV-2 was already significantly inactivated on the TiO_2_ surfaces after 20 min illumination. After 5 h no detectable active virus remained (Figure 1B). Significantly, SARS-CoV-2 on the untreated surface was still fully infectious at 5 h post-addition of virus. TiO_2_–Ag appeared somewhat less effective than TiO_2_ alone, but the difference was not significant.

Plotting the experimental data (Fig. 1B) as ln(titre) v. time (Fig. 1C) yields a disinfection rate coefficient *k* of (5.2 ± 0.6) × 10^−4^ s^−l^, which corresponds to the transport-limited fastest possible rate estimated for SARS-CoV-2 approaching a disinfecting surface in water.^14^ Hence we infer that the viruses arriving at the surface from the inoculum are essentially immediately inactivated. From our illumination conditions we estimate the generation rate of radicals as about 10^13^ cm^−2^ s^−l^,^6^ corresponding to about 800 radicals s^−l^ over the area occupied by one virus at the surface, By extrapolating the data from the first four points to the assumed detection limit, it can be seen that very likely no detectable virus from the initial inoculum remained soon after 2 h exposure (Fig, 1C).

## Discussion

The potent extended anti-SARS-CoV-2 effect of titanium dioxide surface coatings is highly desirable in hospital settings where both patients and staff might be shedding viruses. An important advantage of these surfaces is that they can be activated by ordinary interior light and do not need UV irradiation, which is usually incompatible with simultaneous human presence. The coating has a rough surface with high local curvature that creates an absorption tail into the blue region of the visible spectrum,^6^ overlapping the spectral output of ordinary interior lighting. This is sufficient to ensure an adequate rate of radical generation for effectively immediately inactivating viruses and other microorganisms arriving from the air or hand touches. Conversely, a limitation is that if a sudden very large contamination event occurred, particularly one that severely diminished the light reaching the photocatalyst, it might take impracticably long for the contamination to be eliminated. Hence, in that case rough cleaning, even washing with water, should be used to remove the gross contamination.

The efficacy of the TiO_2_ coating under typical hospital lighting makes it a promising candidate for enhancing the protection afforded by facemasks and other PPE, and well as surfaces likely to be contaminated and hence acting as reservoirs for transmitting infection if left untreated.

## Acknowledgment

The authors thank Saba Yussouf (Invisi Smart Technologies) for helpful discussions. We would also like to thank Nigel Temperton for plasmids.

